# Repetitive magnetic stimulation with iTBS600 induces persistent structural and functional plasticity in mouse organotypic slice cultures

**DOI:** 10.1101/2025.02.23.639712

**Authors:** Han Lu, Shreyash Garg, Maximilian Lenz, Andreas Vlachos

**Affiliations:** Department of Neuroanatomy, Institute of Anatomy and Cell Biology, Faculty of Medicine, University of Freiburg, 79104 Freiburg, Germany; BrainLinks-BrainTools Center, University of Freiburg, 79104 Freiburg, Germany; Center for Basics in Neuromodulation (NeuroModulBasics), Faculty of Medicine, University of Freiburg, 79104 Freiburg, Germany.

**Keywords:** magnetic stimulation, intermittent theta-burst stimulation (iTBS), *in vitro*, Hebbian plasticity, firing rate homeostasis, structural plasticity

## Abstract

**Background:** Repetitive transcranial magnetic stimulation (rTMS) is well known for its ability to induce synaptic plasticity, yet its impact on structural and functional remodeling within stimulated networks remains unclear. This study investigates the cellular and network-level mechanisms of rTMS-induced plasticity using a clinically approved 600-pulse intermittent theta burst stimulation (iTBS600) protocol applied to organotypic brain tissue cultures.

**Methods:** We applied iTBS600 to entorhino-hippocampal organotypic tissue cultures and conducted a 24-hour analysis using c-Fos immunostaining, whole-cell patch-clamp recordings, time-lapse imaging of dendritic spines, and calcium imaging.

**Results:** We observed long-term potentiation (LTP) of excitatory synapses in dentate granule cells, characterized by increased mEPSC frequencies and spine remodeling over time. c-Fos expression in the dentate gyrus was transient and exhibited a clear sensitivity to the orientation of the induced electric field, suggesting a direction-dependent induction of plasticity. Structural remodeling of dendritic spines was temporally linked to enhanced synaptic strength, while spontaneous firing rates remained stable during the early phase in the dentate gyrus, indicating the engagement of homeostatic mechanisms. Despite the widespread electric field generated by rTMS, its effects were spatially and temporally precise, driving Hebbian plasticity and region-specific spine dynamics.

**Conclusions:** These findings provide mechanistic insights into how rTMS-induced LTP promotes targeted plasticity while preserving network stability. Understanding these interactions may help refine stimulation protocols to optimize therapeutic outcomes.

## Introduction

How functional and structural plasticity coordinate on short and long time scales to retain flexibility and stability remains one of the most intriguing questions in neuroscience [1]. From a fundamental neuroscience perspective, this question involves two key dimensions: the forms in which plasticity manifests and the regulatory mechanisms following perturbations in neural activity. Electrophysiological recordings and imaging studies have revealed that neurons adapt to activity perturbation through changes in synaptic transmission and dendritic spine morphology, corresponding to functional and structural plasticity, respectively, on fast and slow time scales [2, 3]. Since the discovery of spine enlargement during long-term potentiation (LTP) [4] and spine shrinkage during long-term depression (LTD) [5] of synaptic transmission, numerous studies have demonstrated the dynamic interaction between these forms of plasticity under basal conditions [6, 7], during learning [8, 9, 10], after pharmacological treatment [11, 12], and in the context of injury or lesion [13, 14], disease [15, 16, 17], or brain stimulation [18, 19]. However, most studies have focused on either functional or structural plasticity in isolation, leaving the interplay between these two phenomena largely unexplored. Furthermore, neural activity, which serves both as the driving force behind activity-dependent plasticity and as a process influenced by the resulting plastic changes, is often insufficiently considered in experimental designs. To address this complex feedback loop, activity perturbation methods are essential. Chronic activity perturbations can be induced through drug administration, pathway lesions, or sensory stimulation and deprivation. Meanwhile, non-invasive brain stimulation (NIBS)—including repetitive magnetic stimulation (rMS)—has gained traction for probing casual relationships in brain function and plasticity, further advancing both fundamental [20] and clinical applications [21].

Repetitive transcranial magnetic stimulation (rTMS) exerts its effects by modulating neural activity [22], altering network dynamics [23], and inducing lasting plastic changes [18, 24]. While numerous studies have demonstrated rTMS-induced functional plasticity—showing that it disrupts the excitation-inhibition (E/I) balance by enhancing excitatory synaptic transmissions [18, 25, 26, 27] and reducing inhibition [24, 28]—its interplay with structural plasticity remains less explored. Structural modifications, such as changes in spine sizes [18], spine turnover dynamics [29], or alterations in dendritic arborization [30] have been reported but are insufficient to elucidate the interplay between rTMS-induced functional and structural plasticity. Moreover, while changes in E/I balance and LTP-induction are often regarded as the primary aftereffects of rTMS [18, 31], their role in driving structural remodeling and influencing network activity dynamics remains less understood. Studies have described transient increases in neural activity after single sessions and enhanced evoked activity following multiple rTMS [32], framing changes in neural activity as a prerequisite for synaptic adjustments. However, this perspective often overlooks the reciprocal interplay between neural activity and plasticity during the stimulation process, which appears central to understanding TMS effects. Recent optogenetic studies tracking neural morphology and activity over extended periods challenge interpretations based solely on Hebbian-like plasticity [33], particularly when simultaneous and long-term data on neural activity and plasticity are lacking.

In this study, we aimed to comprehensively track structural and functional plasticity, as well as network dynamics, over a 24-hours period following a single rMS session. Using entorhino-hippocampal tissue cultures, we applied a 600-pulse iTBS protocol in vitro, which is FDA-approved for treating major depressive disorders [18, 24, 27, 34]. Our findings revealed region-specific plasticity, with prominent expression of the immediate early gene c-Fos in the dentate gyrus and sensitivity to the orientation of the induced electric field. Additionally, an LTP-response of excitatory synapses onto dentate granule cells was observed 2-4 hours and at 24 hours post-stimulation. Time-lapse imaging of neuronal morphology revealed dynamic dendritic spine changes, including enlargement, retraction, and re-enlargement, reflecting a biphasic plasticity response. Calcium imaging revealed rMS-effects at the network level. Despite significant synaptic changes and increased evoked activity in the local network, spontaneous firing rates at the network level remained stable in the dentate gyrus, suggesting the recruitment of homeostatic mechanisms following iTBS-rMS.

## Materials and methods

### Ethics statement

The current study used mouse pups from C57BL/6J (wild type) and Thy1-eGFP mouse lines to prepare entorhinal-hippocampus tissue cultures. All animals were kept under a 12 h − 12 h light-dark cycle with food and water supplied *ad-libitum*. All animal experiments were approved by the appropriate animal welfare committee and the animal welfare officer of Albert-Ludwigs-University Freiburg, Faculty of Medicine under X-17/07K, X-21/01B, and X-17/09C. Special care was taken to ensure the welfare of the animals.

### Preparation of tissue cultures

Entorhinal-hippocampus organotypic tissue cultures (OTCs) were prepared with P3-P5 C57BL/6J or Thy1-eGFP mouse pups as published before in our lab [14, 35]. All tissue cultures were cultured on an insert membrane (Millicell 0.4 *µ*m*/*30 mm, Sigma) in an interface manner for at least 18 days inside the incubator with a humidified atmosphere (5% CO_2_ at 35^◦^C to reach an equilibrium state. The incubation medium consist of 50% (v/v) 1× minimum essential media (#21575 − 022, Thermo Fisher, USA), 25% (v/v) 1× basal medium eagle (#41010 − 026, Thermo Fisher, USA), 25% (v/v) heat-inactivated normal horse serum, 25 mM 1 M HEPES buffer solution (#15630 − 056, Gibco), 0.15% (w/v) sodium bicarbonate (#25080 − 060, Gibco), 0.65% (w/v) glucose (#RNBK3082, Sigma), 0.1 mg*/*ml streptomycin, 100 U*/*ml penicillin, and 2 mM glutamax (#35050 − 061, Gibco). The incubation medium was renewed every Monday, Wednesday, and Friday. If not stated otherwise, the incubation medium applied to tissue cultures was always pre-warmed to 35^◦^C and adjusted around pH = 7.3.

### Repetitive magnetic stimulation

We performed magnetic stimulation with Magstim Rapid^2^ stimulator (Magstim Company, UK). Before magnetic stimulation, tissue cultures were usually transferred to a 35 mm petri dish with 1 ml incubation medium one day in advance. During stimulation, cultures were placed around 1 cm right beneath the coil center. If not stated otherwise, cultures were oriented so that the induced electric field was parallel to the somatodendritic axis of the CA1 pyramidal cells. In a subset of control experiments, we turned cultures 90 degrees counterclockwise so that the induced electric field was perpendicular to the somatodendritic axis of CA1 pyramidal cells. The control cultures were accordingly placed outside the incubator next to the stimulator. The current study used an intermittent theta burst stimulation (iTBS) protocol, which has been approved by the USA Food and Drug Administration (FDA) for treating major depressive disorder (MDD) [34]. 600 pulses were packed as 3-pulse bursts at 50 Hz delivered every 200 ms (5 Hz). Therefore, in each duty cycle of total 20 cycles, 10 bursts were delivered within 2 s and followed by a 8 s interval. All cultures were returned to the incubator after stimulation and revisited for different measurements at certain times. If not otherwise stated, the term *iTBS-rMS* was used throughout the manuscript to refer to magnetic stimulation. We repeated the experiments in two to three rounds in the entire project. Each culture was coded beforehand in a design sheet based on batch number and group information, so that the group information was blind to experimenters during quantification to avoid bias.

### Entorhinal cortex lesion

To apply an entorhinal cortex lesion (ECL), the perforant path was transected from the rhinal to the hippocampal fissure. We performed the ECL 24 hours before magnetic stimulation.

### Immunohistochemical staining

To examine neural activation by magnetic stimulation in tissue cultures, we probed cultures for immediate early gene c-Fos expression as described before [33]. OTCs were fixed with 4% paraformaldehyde (PFA) with 20% sucrose at 90 min and 24 h after iTBS-rMS and stained accordingly. Fixed cultures were first washed three times with 1× PBS (pH = 7.38, 3 × 10 min) to remove residual PFA and then blocked at room temperature (RT) with 10% (v/v) normal goat serum in PBS with 0.5% (v/v) Triton X-100. Later, we incubated the cultures at 4^◦^C with rabbit anti-c-Fos (Synaptic Systems Cat# 226 008, RRID:AB 2891278, 1: 1000) in 1× PBS with 10% (v/v) normal goat serum and 0.05% (v/v) Triton X-100 for 48 h. After incubation with the primary antibody, the cultures were rinsed with 1× PBS (3 × 10 min) and again incubated with Alexa568 anti-rabbit (1: 1000) in 1× PBS with 10% (v/v) normal goat serum and 0.05% (v/v) Triton X-100 overnight at 4^◦^C. In the next morning, cultures were rinsed with 1× PBS (3 × 10 min) and incubated with DAPI (1: 2000) for 20 min at RT. After another 4 washes with 1× PBS (4 × 10 min), we mounted the cultures on glass slides with DAKO anti-fading mounting medium (#S302380-2, Agilent) for confocal microscope imaging.

### Electrophysiological recordings

#### Whole-cell patch-clamp recordings

To further examine whether functional plasticity was triggered in the dentate gyrus, whole-cell patch-clamp recordings were performed on granule cells during 2 to 4 hours after iTBS-rMS and 24 hours after iTBS-rMS. All recordings were conducted at 35^◦^C. The bath solution contained (in mM) 126 NaCl, 2.5 KCl, 26 NaHCO_3_, 1.25 NaH_2_PO_4_, 2 CaCl_2_, 2 MgCl_2_, and 10 glucose (aCSF) and was continuously oxygenated with carbogen (5% CO_2_/95% O_2_). Glass patch pipettes had a tip resistance of 4 − 6 MΩ, filled with the external solution which contained (in mM) 126 K-gluconate, 10 HEPES, 4 KCl, 4 ATP-Mg, 0.3 GTP − Na_2_, 10 PO-Creatine, 0.3% (w/v) biocytin. The external solution was adjusted to pH = 7.25 with KOH and reached 290 mOsm with sucrose). We patched (3-6 neurons per culture) to record the miniature excitatory postsynaptic currents (mEPSCs) of dentate gyrus granule cells in the voltage-clamp mode with a holding potential of −70 mV in the presence of 10 *µ*M D-AP5, 10 *µ*M SR-95531 and 0.5 *µ*M TTX. Series resistance was monitored before and after each recording, and the neuron data was excluded if the series resistance went above 30 MΩ. Each neuron was recorded for around 2 min. Neuron identity was classified based on the resting potential and intrinsic membrane properties.

### Time-lapse imaging

We performed time-lapse imaging in two experiments: (i) tracking the neural dendritic morphology of granule cells before and after stimulation and (ii) monitoring neural calcium activity in the area of the dentate gyrus after stimulation. Live cell imaging was performed at a Zeiss LSM800 microscope with 10× water-immersion (W N-Achroplan 10×/0.3 M27; 420947-9900-000, Carl Zeiss) and 63× water-immersion objectives (W Plan-Apochromat 63×/1,0 M27; 421480-9900-000, Carl Zeiss).

#### Imaging granule cell morphology

To inspect whether the density and size of the neural spine were associated with altered functional transmission among granule cells within 24 h after iTBS-rMS, we employed the time-lapse imaging technique to follow the same apical dendritic segments of granule cells several days before and after stimulation. Thy1-eGFP tissue cultures were used in which clear granule cells could be identified. To characterize baseline and normal fluctuations, we imaged the same dendritic segment every day at the same time for 5 days from *day 0* to *day 4*. On *day 4*, we imaged the dendritic segment prior to magnetic stimulation and performed iTBS-rMS on tissue cultures ca. 1 h later. The same dendritic segment was then imaged every hour for 5 hours after iTBS-rMS and again on *day 5*, that is, at 24 h after iTBS-rMS. The imaging protocol was performed as previously described in [35].

#### Calcium imaging

Calcium imaging was performed in tissue cultures prepared from wild-type animals to further inspect whether magnetic stimulation changed the activity of the neural network within the dentate gyrus. The procedure was previously described in another study [36]. Tissue cultures (between DIV3 and DIV5) were transfected with 1 *µ*L AAV1-hSyn1-GCaMP6f-P2A-nls-dTomato virus (Addgene viral prep #51085-AAV1, http://n2t.net/addgene:51085, RRID:Addgene 51085, a gift from Jonathan Ting) diluted 1: 4 in 1× PBS (pH = 7.38), by pipetting a drop of the mixture on top of the culture to cover the entire culture. The virus will express the calcium indicator GCaMP6f and the labelling fluorescent protein TdTomato in the transfected neurons. To approximate the network dynamics of the tissue cultures inside the incubator, we used a pre-warmed incubation medium with pH adjusted to 7.38 throughout the imaging session for calcium signal. During the imaging session, the membrane insert with 2 cultures was placed into a 35 mm petri dish filled with 5 ml incubation medium. We imaged three batches of cultures; each batch contained cultures recorded 3 hours after iTBS-rMS and 24 hours after iTBS-rMS and their corresponding controls.

### Confocal microscope imaging for fixed samples

A Leica SP8 laser-scanning microscope acquired fluorescent images of DAPI and c-fos in the fixed tissue. We used the 20× multi-immersion (NA 0.75; Leica) objective to take tile-scan the whole culture stack at 512 × 512 px resolution with Δ*z* = 2*µ*m. The laser intensity was adjusted accordingly to achieve a comparable non-saturated fluorescence intensity among all groups.

### Quantifying microscope images

#### c-Fos positive cell counting and normalisation

c-Fos signals were quantified by the segmentation and counting functions of Fiji ImageJ. To ensure a comparable baseline across cultures with variable thickness, we projected the five middle planes of c-Fos and DAPI *z*-stacked images with a maximum intensity. Four regions of interest (ROIs), including EC, DG, CA3, and CA1, were drawn based on each culture’s cell aggregation pattern in the DAPI signal. These masks were applied to the same culture’s c-Fos signal for further analysis of each subregion. We used macro that turned the images into binary, segmented the cells based on size and circulatory, and then returned the cell count of each ROI. The Fiji macro script is included in the supplemental materials repository. To pool several batches and avoid the bias of varied expression baselines, we normalised each group’s c-Fos positive cell count in each batch by the corresponding mean of the control group.

#### Spine density counting and normalisation

*z*-stacked fluorescent images of Thy1-eGFP cultures were projected to create a 2*D* representation of individual dendritic segments. ImageJ plugin Spine Density Counter [37] was used to count spine numbers and measure segment length, which estimates spine density. Special attention was paid to the same dendritic segments imaged at different times to ensure that the same starting and ending points were used. *Post hoc* visual inspection was conducted to ensure that the spine detection results were not strongly biased. To visually differentiate the potential changes, we normalised the spine density of each segment by dividing it by its baseline spine density at *day 4* sampled right before stimulation.

#### Spine size estimation, normalisation, and binning

Spine size quantification and analysis were performed as described before [35]. The same *z*-projected fluorescent images were used and converted to binary to track individual spines for spine size analysis. To eliminate bias from drawing and automatic reconstruction, we drew circles manually around the spine to cut it from the dendrite; the spine size was estimated by measuring the signal intensity with an arbitrary unit of the drawn circle. The drawing and measurements were performed with FIJI ImageJ. Out of the same principle, we also normalised the time series of individual spine size over 11 time points by dividing the corresponding size values by its size at *day 4* right before any stimulation and frequent imaging sessions. In the analysis, both raw and normalised spine size values were used.

In the following analysis, we normalised the alteration of spine sizes (Δspine size) by their size at *day 4* and binned the data by their initial sizes. We first fetched the spine sizes of individual spines at *day 4* and a given timing (for instance, 2 h, 5 h, or 24 h after the iTBS-rMS were used) and calculated the alteration (Δspine size) by the size differences between *day*4 and the given time. Then, we normalised the amount of alteration by the corresponding spine size at *day 4*. In the third step, we sorted and binned the spines by their initial sizes at *day 4*. By collecting spine IDs in each binned group, we grouped the spine size alterations (Δspine size) by their initial spine size at *day 4*. This analysis shows us how spines with different initial sizes update their size over a certain time course with and without perturbation.

#### Analyzing calcium imaging data

We used a similar computer vision algorithm to create segmentation masks based on TdTomato signals as described before [36]. The TdTomato images were pre-processed to retain either the dentate gyrus blade area or the hilus area. Two sets of masks were applied to segment and extract the intensity of the eGFP signal of the calcium indicator GCaMP6f frame by frame in the DG and the hilus area, respectively.

The raw trace was detrended by subtracting the median value with a rolling method (window size: 20) and then normalized based on the tread mean in each trace to achieve Δ*F/F*_0_ traces. Calcium spikes were automatically detected for each processed trace with a threshold of 3 times the standard deviation from the mean values. Individual traces and spike detection data were visually inspected for quality control. In addition to calcium spikes, we used the *trapz* function in *Python* to sum up the area under the curve to account for subthreshold dynamics.

### Analyzing electrophysiological recordings

#### mEPSCs analysis

Excitatory postsynaptic currents were analyzed using the automated event detection tool from the pClamp11 software package as previously described [14, 24, 36]. For plotting the average distribution of mEPSC amplitudes and areas, we calculated the distribution of event amplitudes/areas of individual neurons and then averaged all distributions for each group. To obtain the summated area of each neuron over 2 minutes, we first summated the total event areas over the entire recording period. Since the recording period is not exactly 2 minutes in practice, we scaled the value by an coefficient (2 minutes over the actual recording period in minutes as a unit).

### Statistical analysis

We showed the error bars with s.e.m. values in all the figures. All experiments were repeated in two to three independent batches. Statistical significance was assessed with Mann-Whitney U tests (c-Fos data), two-way analysis of variance (ANOVA; c-fos data in ECL experiments), One-way ANOVA (mEPSCs events), Two-way RM ANOVA (time series of spine density and spine sizes), Kruskal-Wallis test (correlation coefficient among calcium activity) or linear mixed models (LMM) where data clustering occurred within each neuron (mEPSCs events) or each culture (calcium spikes, calcium trajectory AUCs, and power spectra AUCs). For multiple testing, *p* values were adjusted accordingly with Tukey’s multiple comparison test (for One-way ANOVA), Dunn’s multiple comparison test (for Kruskal-Wallis test), or Sidak’s multiple comparison test (for Two-way RM ANOVA). Statistical analysis was performed using Prism 9.0 GraphPad or *Python* package *statsmodels.formula.api* (for linear mixed model analysis). The statistical test, *p* value, and the group sizes were included in the text and the corresponding figure legends. If not stated otherwise, ∗ is *p <* 0.05, ∗∗ is *p <* 0.01, ∗ ∗ ∗ is *p <* 0.005, while “ns” means *p* ≥ 0.05. The digits at the bottom of each bar label each group’s culture or neuron numbers. For LMM analysis, we reported the confidential interval (CI) of the estimated coefficient. The effects were insignificant if 95%CI crosses zero or significant if 95%CI, 99%CI, 99.95%CI or even 99.99%CI does not cross zero.

### Data availability

All datasets and analysis code generated within this project is available in https://github.com/ErbB4/rMS-iTBS.

## Results

### iTBS-rMS induces c-Fos expression in the dentate gyrus

To map regions of plasticity induction in entorhino-hippocampal tissue cultures subjected to iTBS-rMS, we quantified the expression of c-Fos, a well-established immediate early gene and reliable marker of activity-dependent synaptic plasticity. Brain tissue cultures (*>* 18 days *in vitro*) prepared from wild-type mice of both sexes were subjected to a single 600-pulse iTBS protocol, while control cultures were placed alongside stimulated cultures without receiving stimulation (sham control). Tissue cultures were fixed and stained for c-Fos and DAPI at 1.5 h and 24 h post-stimulation (Figure 1A-C). At 1.5 h, we observed a significant increase in c-Fos expression (Figure 1D), in the dentate gyrus (DG) compared to sham controls (*p* = 0.0002, Mann-Whitney U test). In contrast, while the CA3 and CA1 regions showed increased c-Fos levels, these changes were not statistically significant (*p* = 0.0881 and *p* = 0.065, respectively, Mann-Whitney U test). By 24 h post-stimulation, c-Fos levels in all subregions had returned to baseline (*p* = 0.403, *p* = 0.566, *p* = 0.986, respectively for DG, CA3, CA1; Mann-Whitney U test). This transient pattern suggests that iTBS-rMS induced robust c-Fos expression in the DG of our preparations. Interestingly, rotating the cultures to 90 degrees counterclockwise abolished c-Fos upregulation in DG (*p* = 0.533; CA3, *p* = 0.533; CA1 *p* = 0.758; Mann-Whitney U test), indicating that DG activation by iTBS-rMS is orientation-dependent.

**Figure 1:**
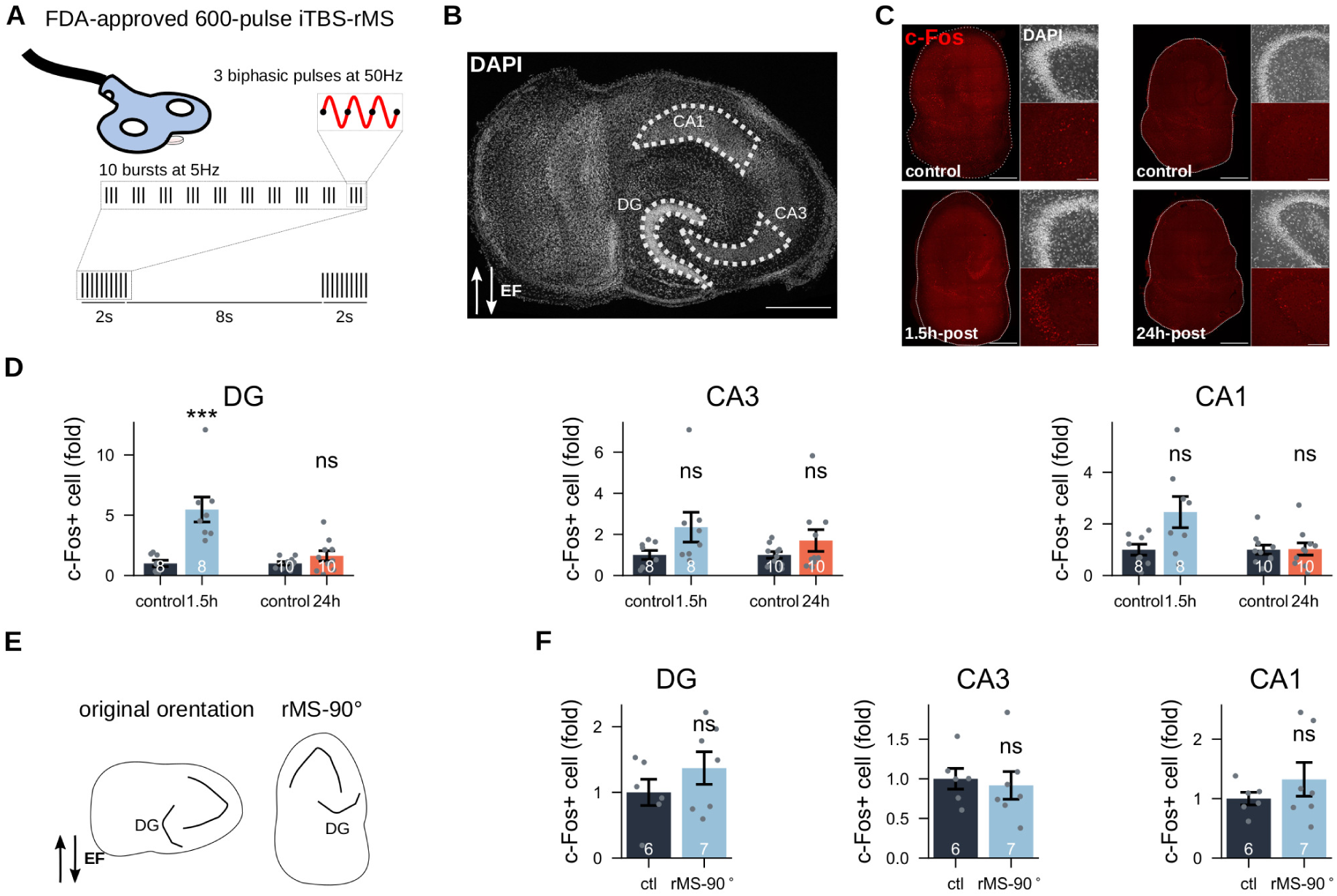
iTBS-rMS induces c-Fos upregulation in the dentate gyrus (DG) in an orientation-dependent manner. (A) The FDA-approved 600-pulses iTBS protocol was applied to three-weeks old entorhino-hippocampal tissue cultures. (B) Segmentation masks delineating the three subregions (DG, CA3, CA1) based on DAPI nuclear staining (Scale bar: 500 *µ*m). (C) Representative c-Fos staining in cultures across different conditions: 1.5 h post-stimulation and corresponding controls; 24 h post-stimulation and corresponding controls (scale bar: 500 *µ*m). Insets show DAPI staining and c-Fos signals in the DG (scale bar: 100 *µ*m). (D) Normalised c-Fos-positive cell counts for each group across the three subregions. Statistical analysis was performed using the Mann-Whitney U test; unless otherwise specified, ∗ is *p <* 0.05, ∗∗ is *p <* 0.01, ∗ ∗ ∗ is *p <* 0.005, while “ns” means *p* ≥ 0.05. The number below each bar represents the sample size (=cultures) per group. Bars and error bars denote the mean and standard error of the mean (s.e.m), respectively. (E) Schematic illustration of magnetic stimulation following a 90 degrees rotation of the cultures. (G) Normalised c-Fos positive cell counts in control cultures and those stimulated after rotation.

### iTBS-rMS enhances excitatory transmission in granule cells

To investigate the link between c-Fos upregulation and activity-dependent plasticity in granule cells, we conducted whole-cell patch-clamp recordings of dentate granule cells in the suprapyramidal blade of the DG. AMPA receptor-mediated miniature excitatory postsynaptic currents (mEPSCs) were recorded across three groups: sham controls, cultures assessed 2 − 4 h after iTBS-rMS, and cultures analyzed 24 h post-iTBS-rMS (Figure 2A). Analysis of mEPSC amplitude distributions (Figure 2B) revealed a significant increase in the 2−4 h post-stimulation group, with no similar shift observed at 24 h post-stimulation (*p <* 0.0001, One-way ANOVA with Tukey’s multiple comparison test), findings corroborated by linear mixed model (LMM) analysis (99.99%CI = [−6.476, −1.235]). A comparable increase was observed in the averaged distributions of mEPSCs areas (Figure 2C; *p <* 0.0001, One-way ANOVA with Tukey’s multiple comparison test; 95%CI = [−0.384, 1.130], LMM), indicating an enhanced synaptic efficacy shortly after stimulation. mEPSCs frequencies significantly increased in dentate granule cells recorded both 2−4 h and 24 h post-stimulation compared to the sham controls (Figure 2C; *p* = 0.0036 for 2−4 h-post, *p* = 0.0008 for 24 h-post, Kruskal-Wallis test with Dunn’s multiple comparison test), with no significant differences between the two post-stimulation time points (*p >* 0.9999, Kruskal-Wallis test with Dunn’s multiple comparisons test). Quantification of overall synaptic transmission, based on aggregate synaptic event areas over a 2-minute period, revealed significant increases at both 2−4 h (*p* = 0.0013, Kruskal-Wallis test with Dunn’s multiple comparison test) and 24 h post-stimulation (*p* = 0.0031, Kruskal-Wallis test with Dunn’s multiple comparison test; Figure 2D), suggesting a lasting potentiation of excitatory input to granule cells. No significant differences in synaptic strength were observed between the early and the late post-stimulation phases (*p >* 0.9999, Kruskal-Wallis test with Dunn’s multiple comparison test). These results demonstrate that iTBS-rMS promotes a rapid and transient peak in synaptic strength within hours after stimulation, followed by persistent enhancement lasting up to 24 h post-stimulation.

**Figure 2:**
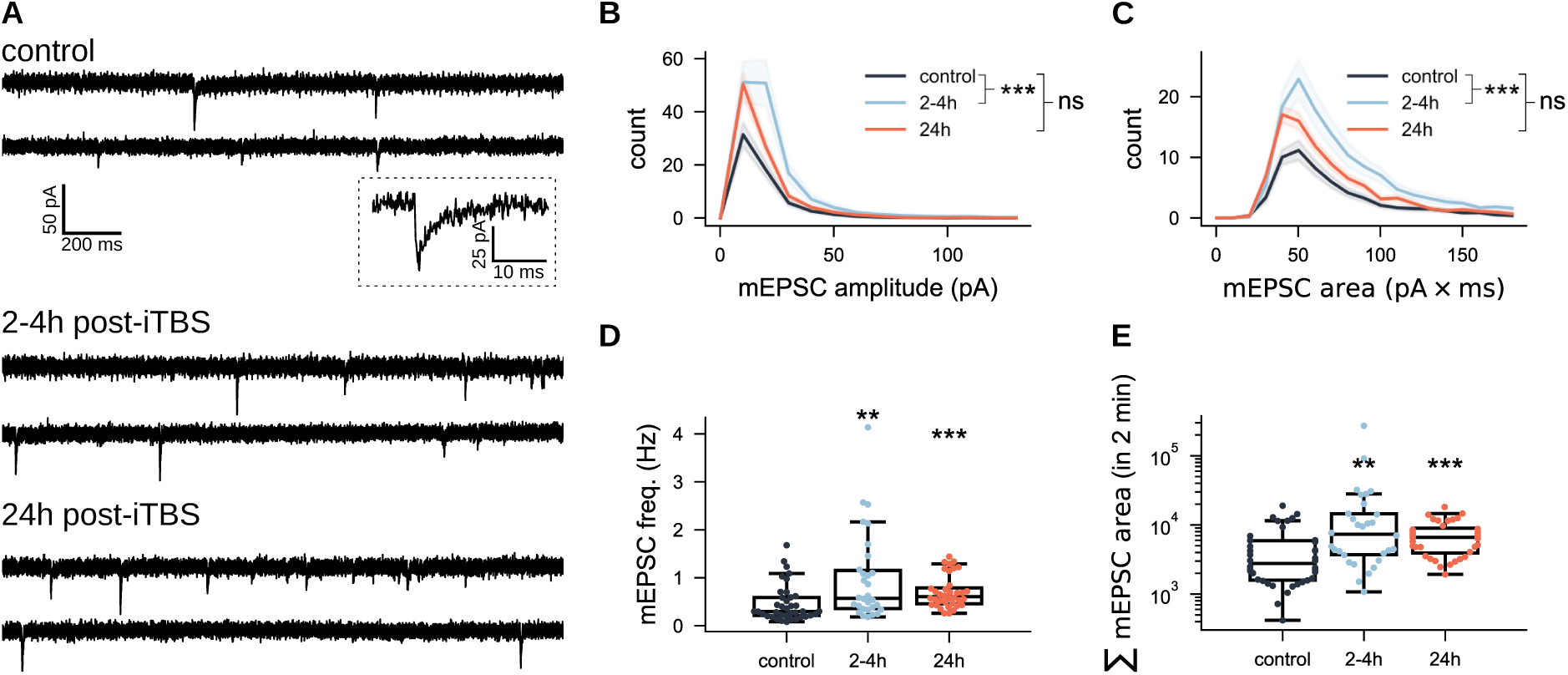
iTBS-rMS enhances synaptic strength in DG granule cells. (A) Representative sample traces of AMPA receptor-mediated miniature excitatory postsynaptic currents (mEPSCs) recorded from dentate granule cells in the three experimental groups (control, 2-4 hours, and 24 hours post-iTBS600-rMS). (B,C) Average distributions of mEPSC amplitudes and areas across recorded neurons for each group. A total of *N* = 36 neurons were recorded from control cultures, *N* = 30 neurons from cultures 2 h-4 h post-stimulation, and *N* = 36 neurons from cultures 24 hours post-stimulation. As no significant differences were observed 2−4 h and 24 h control groups, data from both control groups were pooled. (D) mEPSC frequencies across the three groups. Each dot represents the frequency of a recorded neuron. (E) Overall synaptic strength in the three groups, quantified by summing mEPSC areas over a standardized 2-minute recording duration. Each dot represents an individual neuron.

### iTBS-rMS modulates dendritic spine morphology in granule cells

To examine whether the functional changes in synaptic transmission correspond to morphological changes of dendritic spines, we performed time-lapse imaging experiments in tissue cultures prepared from Thy1-eGFP mice, where principal neurons are visualized by enhanced green fluorescent proteins (eGFPs; Figure 3A). Dendritic segments were imaged daily over a 5-day baseline period, with iTBS-rMS applied on *day 4*, approximately 1 h after the final baseline imaging session. Post-stimulation, cultures were imaged hourly for 5 hours and once more 24 h later to capture both rapid and delayed morphological changes (Figure 3B,C).

**Figure 3:**
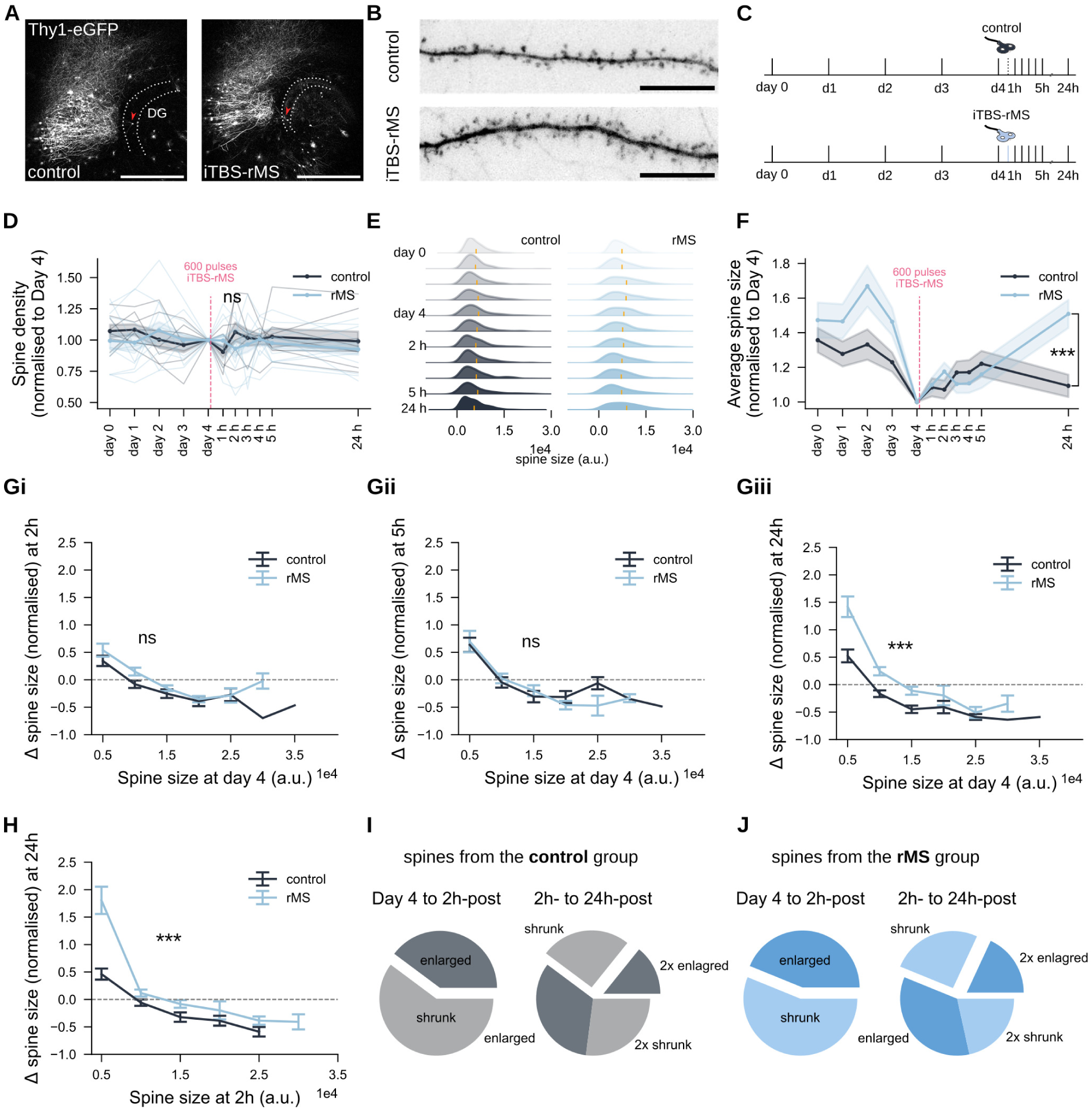
iTBS-rMS modulates spine sizes but not densities. (A) Time-lapse imaging of dentate granule cells in tissue cultures from Thy1-eGFP mice was performed to track dendritic spines and their morphologies in control and iTBS600-rMS groups. Red arrows indicate example granule cells shown in (B). (Scale bar: 500 *µ*m.) (B) Representative dendritic segments from both groups. (Scale bar: 10 *µ*m.) (C) Schematic representation of the experiment protocol. (D) Average spine density in both groups, normalised to the respective spine density on *day 4* for each segment. (*N* = 6 control, *N* = 5 stimulated cultures, from three independent experiments.) (E) Pooled distributions of spine sizes across 11 time points. (*N* = 211 spines in controls, *N* = 260 spines in the stimulated group) (F) Mean normalised spine sizes with standard error of the mean (s.e.m.) for both groups. (Gi-Giii) Normalised changes in spine sizes at 2 h, 5 h, or 24 h post-stimulation, grouped by their baseline spine size on *day 4* prior to the stimulation. The *x*-axis values represent the upper limits of each binning group. Data points above the dashed line indicate spine enlargement, while those below indicate spine shrinkage. (I,J) Proportion of spines that enlarged or shrank at 2 h post-stimulation relative to *day 4*, and at 24 h post-stimulation relative to to 2 h post-stimulation.

Spine density analysis revealed fluctuations during the 5-day baseline period, with no significant changes following iTBS-rMS at any time point (*p >* 0.99, Two-way RM ANOVA with Sidak’s multiple comparison test; Figure 3D). Spine size analysis showed a long-tail distribution throughout the imaging sessions, with most spines being small and a few being large (Figure 3E). Notably, iTBS-rMS broadened the size distribution at 24 h post-stimulation, characterized by a significant increase in mean spine size (*p* = 0.0005, Two-way RM ANOVA with Sidak’s multiple comparison test; (Figure 3F).

To investigate size-dependent dynamics, changes in spine size (Δ spine size) were analyzed at 2 h, 5 h, and 24 h post-stimulation, normalised to baseline sizes on *day 4* right before iTBS-rMS (Figure 3Gi-Giii). In control cultures, small spines tended to enlarge while large spines shrank. iTBS-rMS enhanced these dynamics, with small spines enlarging further and large spines shrinking less than in controls. While the enhancement was insignificant at 2 h (*p* = 0.165, Unpaired t-test) and diminished at 5 h (*p* = 0.5657, Unpaired t-test), it became pronounced again at 24 h (panel Giii; *p* = 0.0001, Unpaired t-test). These results align with mEPSC data, supporting the link between functional and structural synaptic potentiation observed at similar time points after stimulation.

To determine whether the spines enlarged at 24 h originated from those already enlarged at 2 h or from a different subpopulation, we performed a comparative analysis of spine sizes dynamics between the two time points (Figure 3H). A similar pattern of enlargement, primarily among small spines was observed (*p* = 0.0007, Unpaired t-test). Further classification of spines into “enlarged” or “shrunk” groups revealed that most spines enlangered at 24 h originated from the group that had *shrunk* at 2 h, rather than from spines already enlarged earlier (Figure 3I).

In the stimulated group, a slightly higher proportion of initially shrunk spines (2 h-post) transitioned to enlargement at 24 h compared to controls, though this trend did not reach statistical significance (Figure 3J; *p* = 0.1866, Chi-square test). We concluded that iTBS-rMS does not significantly alter dendritic spine densities but induces dynamic changes in spine sizes. Small spines show early enlargement at 2 h post-stimulation, while a pronounced enlargement occurs at 24 h, driven by distinct spine subpopulations. These findings build upon and expand our earlier results from the CA1 region in response to 10 Hz stimulation [18], emphasizing the temporal complexity of iTBS-induced synaptic remodeling. They suggest that functional potentiation is intricately linked to morphological adaptations within specific dendritic spine populations.

### iTBS-rMS modulates spine size dynamics in granule cells

The method used to track spine sizes by assigning spine IDs, inherently incorporates spatial information about spine location and proximity—spines with adjacent IDs were physically closer. This approach enabled an assessment of the effects of iTBS-rMS on heterogeneous structural plasticity under baseline conditions. As shown in Figure 4A, over a closely monitored 24 hour period, each spine exhibited a distinct evolution trajectory, resulting in unique correlation coefficient matrices for each dendritic segment (Figure 4B). To compare matrices of varying dimensions, which were determined by the number of identified spines, unit matrices were generated by averaging a 15 × 15 matrix along its diagonal. Analysis of variance among these unit matrices revealed distinct differences between the control and stimulated segments (Figure 4C, D). In control segments, size changes of proximate spines were positively correlated while distant spines showed negative correlations. iTBS-rMS disrupted this spatial pattern, resulting in uniformly positive correlations among all spines, regardless of their relative distance.

**Figure 4:**
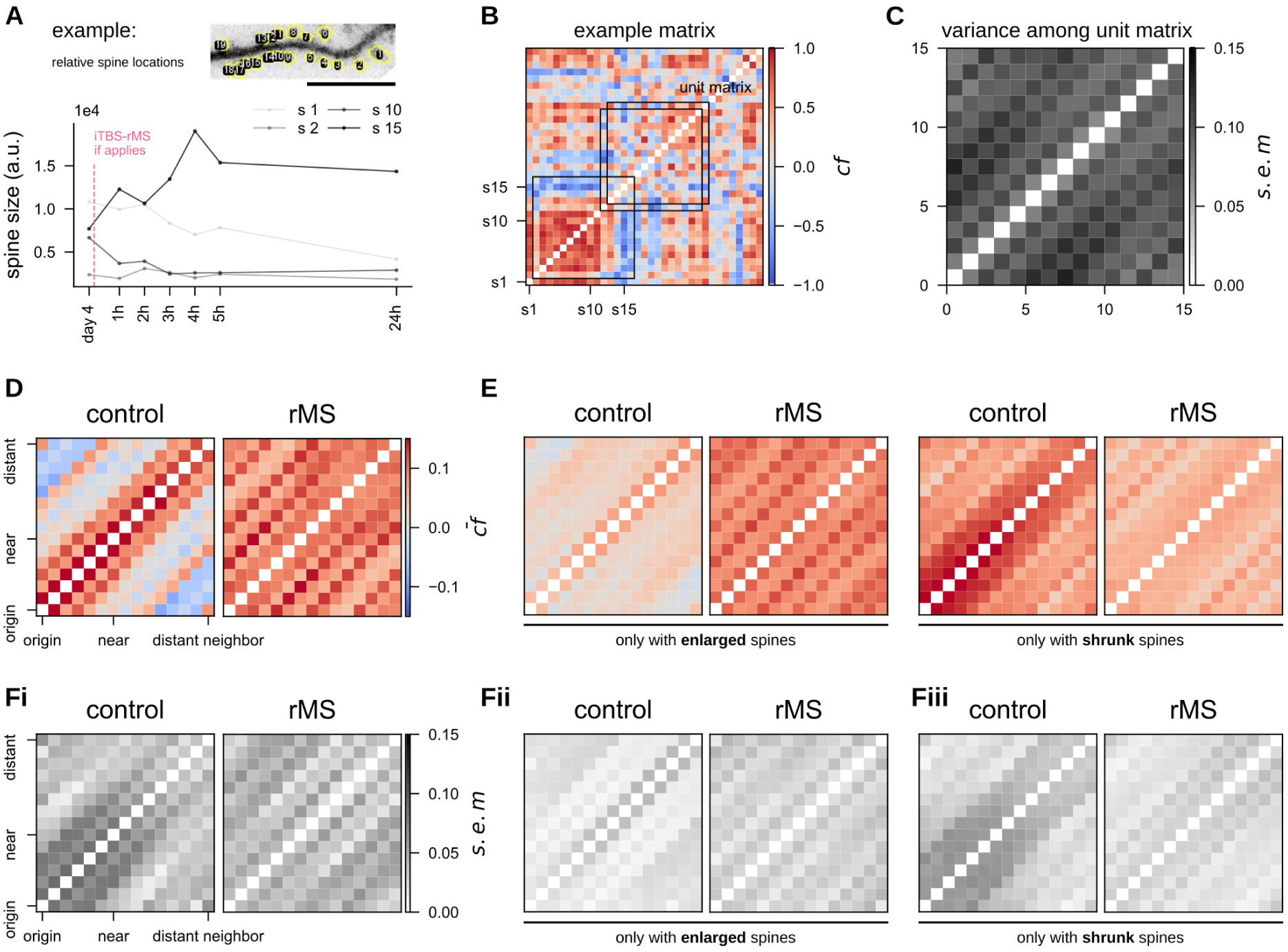
iTBS-rMS modulates heterosynaptic structural plasticity. (A) Representative dendritic segment of a GFP-expressing dentate granule cell illustrating individual spines identified and measured for size changes. (Scale bar: 10 *µ*m.) The four trajectories depict size fluctuations of selected spines over a 24-hour time window. (B) Correlation coefficient matrix of the sample dendritic segment, showing the relationship between any pair of spines. A moving average along the diagonal defines a 15 × 15 unit matrix. (C) Variance among unit matrices obtained by averaging the matrices in (B). (D) Averaged unit matrices for control and stimulated segments. Positive correlations indicate coordinated size changes, while negative correlations suggest opposing alterations in spine size. (E) Averaged unit matrix after classifying spines into enlarged or shrunken groups, based on their size differences between *day 4* and 24 h post-stimulation. (Fi,Fii) Variance among segments when averaging unit matrices across different dendritic segments.

To test whether these changes were determined by spine enlargement following stimulation, spines were categorized as either “enlarged” or “shrunk” at 24 h post-stimulation relative to their baseline size at *day 4*. In the control group, this categorization disrupted the unit matrix pattern (Figure 4D, E), while in the iTBS-rMS group, the unit matrix pattern was largely preserved within the pool of enlarged spines. This preservation likely reflected the pronounced enlargement of *small* spines in response to stimulation. The variance analysis of the unit matrices in individual slice cultures confirmed these observations and highlighted the robustness of the effects of iTBS-rMS (Figure 4Fi-Fii). These findings suggest that iTBS-rMS fundamentally alters the regulation of spine sizes, overriding the natural spatial correlations observed under baseline conditions.

### iTBS-rMS alters neural activity and calcium dynamics in the dentate gyrus and hilar region

Given that whole-cell patch-clamp recordings and time-lapse imaging confirmed potentiation of excitatory synapses among granule cells following iTBS-rMS, we examined whether local network activity was altered. Calcium imaging was performed in the DG, and neuronal time series were extracted using computer vision techniques (Figure 5A). Calcium signal trajectories (Δ*F/F*_0_) of control cultures and those measured 3 h and 24 h hours post-stimulation showed distinct dynamics (Figure 5B). Calcium spikes and the area under the curve (AUC) were calculated to assess both suprathreshold and subthreshold activity (Figure 5C). We observed a significant increase in calcium spikes in neurons from stimulated cultures at 24 h post-stimulation (*p <* 0.0001, Unpaired t-test; 99.95%CI = [0.055, 0.728], LMM) but not 3 h (Figure 5D; *p* = 0.1958, student’s t-test; 95%CI = [−0.178, 0.185], LMM). Similarly, the AUC significantly increased at 24 h, reflecting elevated calcium activity in the DG after the stimulation (Figure 5E; *p* = 0.2735 and *p* = 0.1946 for 3 h and 24 h-post respectively, Unpaired t-test; 95%CI = [−0.398, 1.063] and 95%CI = [−0.304, 1.479], LMM). Despite the increased activity, network synchronization, measured as correlation among individual neurons, did not show significant changes (Figure 5F, *p* = 0.4213 for 3 h, *p* = 0.0643 for 24 h, Mann-Whitney U test). Power spectrum analysis of individual neurons revealed higher power at low frequencies (Figure 5Gi), but group-level analysis of power spectra at both 3 h and 24 h yielded no significant difference after accounting for clustering effects (95%CI = [−2.496, 4.516] and 95%CI = [−1.653, 15.256], LMM). Together, these results indicate that calcium spikes in dentate granule cells increased significantly at 24 h but not 3 h post-stimulation, with no corresponding changes in network synchronization or power spectra.

**Figure 5:**
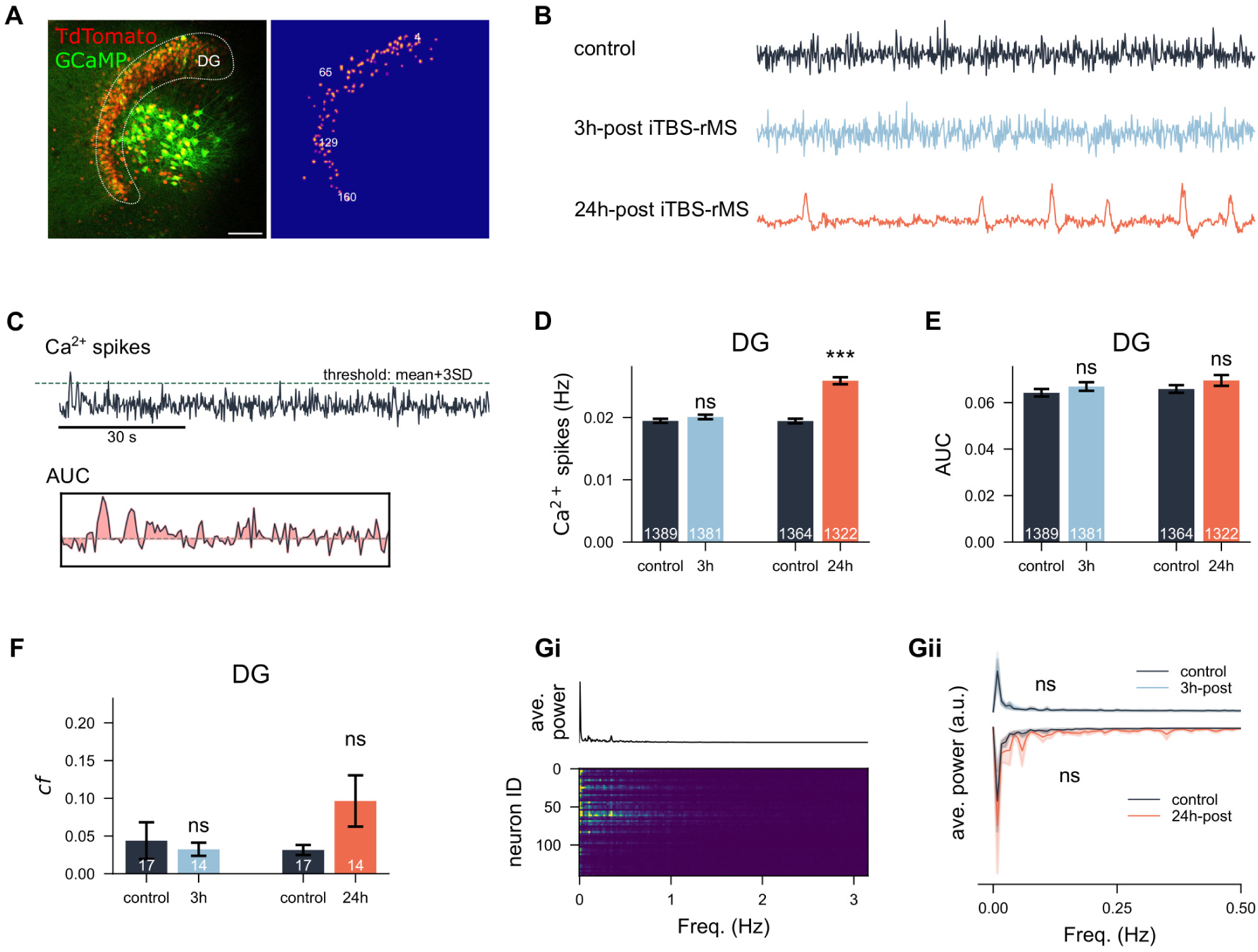
iTBS-rMS enhances dentate gyrus calcium activity 24 h post-stimulation. (A) Calcium imaging of the dentate gyrus (DG). Neurons were transfected with the viral vector AAV1-hSyn1-GCaMP6f-P2A-nls-dTomato. A computer vision algorithm segmented neurons within the DG using a mask based on the TdTomato signal, allowing extraction of the corresponding calcium time series. (Scale bar: 50 *µ*m.) (B) Representative calcium signal trajectories in control cultures and in cultures at 3 h or 24 h post-stimulation. (C) Methods for estimating calcium activity: detection of calcium spikes via thresholding and calculation of the area under the curve (AUC) for each trace. The AUC epoch panel magnifies a 30-second segment shown in the upper panel. (D,E) Mean and standard error of the mean (s.e.m.) of calcium spike frequency and AUC values across four experimental groups. (F) Mean and s.e.m. of correlation coefficients among all recorded calcium traces within each culture, compared across four groups. (Gi) Lower panel: power spectra of individual neurons from a representative culture. Upper panel: averaged power spectra of the sampled culture. (Gii) Average power spectra of each experimental group (3 h or 24 h after stimulation) and their corresponding control cultures. The 24 h group and controls are inverted for visualization purposes only.

A similar analysis was conducted in the hilar region using a modified mask (Figure 6A). Unlike DG neurons, hilus neurons displayed significantly enhanced calcium spikes at both 3 h and 24 h post-stimulation (Figure 6B; *p <* 0.0001 and *p <* 0.0001 for both timings, Unpaired t-test; 99.9%CI = [0.249, 1.187] and 99.9%CI = [0.535, 1.617], LMM). AUC values supported these findings (*p <* 0.0001 and *p* = 0.0003 for both time points, Unpaired t-test; 99.5%CI = [0.450, 6.375] and 95%CI = [0.641, 5.992] for both timings, LMM). Furthermore, the correlation among hilus neurons significantly increased at both time points (Figure 6D; *p* = 0.0101 and *p* = 0.0172, Mann-Whitney U test). While group-level analysis of power spectra indicated significant differences (*p <* 0.0001 and *p* = 0.0316, Unpaired t-test), no significant changes were observed when accounting for clustering effects with LMM analysis (Figure 6E,F; 99.5%CI = [−55.008, 143.532] and 99.9%CI = [−51.786, 80.295] for both timings). These results show that hilus neurons were significantly more active at 3 h and 24 h post-stimulation compared to controls, highlighting region-specific differences in the response of iTBS-rMS.

**Figure 6:**
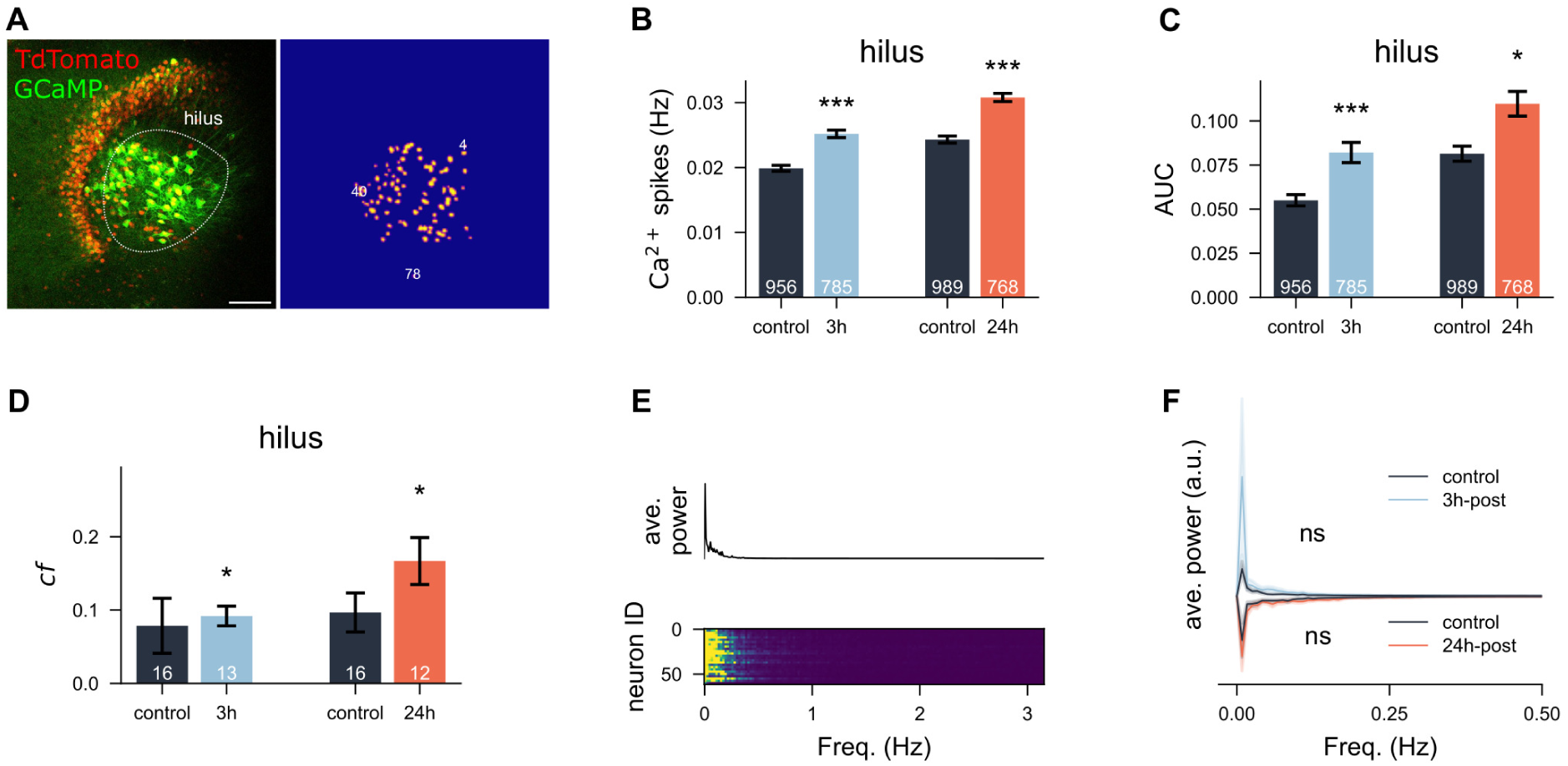
iTBS-rMS increases calcium activity in the hilus at 3 h and 24 h post-stimulation. (A) Representative culture and corresponding segmentation mask for identifying hilus neurons. (Scale bar: 50 *µ*m) (B,C) Mean and standard error of the mean (s.e.m.) of calcium spike frequency and area under the curve (AUC) values across all neurons in four experimental groups. (D) Mean and s.e.m. of correlation coefficients among all recorded calcium traces within each culture, compared across four groups. (E) Lower panel: power spectra of individual neurons from a representative culture. Upper panel: averaged power spectra of the sampled culture. (F) Average power spectra for all four experimental groups.

## Discussion

This study investigated the interplay between functional and structural plasticity, as well as network dynamics within 24 hours following a single session of FDA-approved 600-pulse iTBS magnetic stimulation. We observed that c-Fos expression in the DG was sensitive to the orientation of the induced electric field, indicating that iTBS effectively triggered plasticity induction in a direction-dependent manner. Functionally, synaptic strength was initially enhanced, as reflected by increases in both mEPSC amplitudes and frequencies, alongside spine remodeling over time. Structurally, significant changes involved the enlargement of distinct cohorts of small dendritic spines, which contributed to increased synaptic efficacy. Despite these functional and morphological modifications, calcium imaging revealed no detectable changes in spontaneous network activity in the DG during the early phase after stimulation, while activity levels increased in the hilus. By 24 hours, both synaptic strength and calcium activity in the DG were elevated. These findings highlight the spatial and temporal specificity of iTBS-induced plasticity, suggesting a fine-tuned interaction between Hebbian plasticity and homeostatic regulation.

Hebbian plasticity and firing rate homeostasis have long been considered distinct yet complementary mechanisms essential for maintaining neural network function. While computational studies proposed that their interplay is critical for adaptive network dynamics [38, 39, 40], experimental studies often focus on one mechanism at the expense of the other due to design constraints. By integrating functional, structural, and network-level analysis, our study provides empirical evidence for their coexistence, demonstrating that iTBS-induced Hebbian plasticity, reflected by synaptic strengthening and structural remodeling, occurs alongside firing rate homeostasis. Despite progressive synaptic potentiation, spontenous network activity in the DG remained stable, indicating a compensatory regulation that prevents excessive excitation. This balance between plasticity-driven flexibility and homeostatic stability highlights the intricate coordination required to maintain robust network function following external perturbations and may ultimately shape the long-term efficacy of iTBS600 interventions.

Prior work from our lab identified a persisting increase in mEPSC frequencies in CA1 pyramidal neurons at 6-8 hours post-stimulation [18, 27, 41]. The present results align with and extend these findings, demonstrating that similar persistent increases in mEPSC frequencies occur in DG granule cells and persist up to 24 hours post-stimulation. These observations suggest that the previously described mechanisms of plasticity in CA1 may not be limited to this region but extend well into the DG. Similarly, the preferential increase in the size of small spines observed in our previous work in CA1 is also evident in the DG, highlighting a consistent mechanism of structural plasticity across these brain regions. Importantly, similar to CA1, we did not observe significant changes in spine densities in the DG, suggesting that structural remodeling is driven by changes in spine morphology rather than turnover. The orientation-dependent effects on c-Fos expression observed in our study underscore the spatial specificity of rTMS-induced plasticity. When the orientation of the electric field was altered, c-Fos upregulation in the DG was abolished, suggesting that the spatial alignment of the stimulation plays a crucial role in the activation of plasticity-related pathways. This highlights the importance of considering stimulation geometry in both experimental and clinical applications of rTMS. Whether and how the initial c-Fos expression is linked to these observations remains unknown at this point.

Time-lapse imaging experiments provided critical insight into the temporal coordination of structural plasticity. Under control conditions, dendritic spines exhibited natural size fluctuations, with small spines tending to enlarge and large spines tending to shrink over time, consistent with previously reported homeostatic dynamics [35]. Following iTBS-rMS, these natural dynamics were disrupted, with small spines showing significant enlargement 3 hours post-stimulation. This enlargement diminished by 5 hours, followed by a re-emergence of spine growth at 24 hours post-stimulation. These observations reveal temporally distinct structural plasticity events that align with functional changes observed in mEPSC events.

Transient enlargement of the spine at 2 hours after stimulation may correspond to the potentiation of functional synapses or the unsilencing of previously silent spine synapses, as reflected by the concurring increase in mEPSC amplitudes and frequencies during this period. The subsequent reduction in spine enlargement at 5 h-post could indicate a natural decay or homeostatic adjustment of potentiated synapses similar to the depotentiation observed in CA1 neurons at 6-8 hours post-stimulation. The re-emergence of spine enlargement at 24 hours post-stimulation appears to involve the recruitment of distinct cohorts of spines, as newly enlarged spines at this time point originated primarily from previously unengaged small spines. This supports the idea of cohort-specific recruitment, with different populations of spines contributing to synaptic transmission over time.

In addition to these temporal dynamics, our analysis suggests that structural plasticity during the later phases disrupts baseline spatial correlations in dendritic segments. The enlargement of small spines, which may have been randomly distributed along the dendritic arbor, disrupts the spatially constrained heterosynaptic structural plasticity observed under baseline conditions. This reorganization of the dendritic spine turnover dynamics could reflect a more general adaptive response to iTBS, in which functional and structural plasticity converge to support long-term changes in excitatory input.

Local network activity plays a pivotal role in shaping synaptic and structural plasticity, yet its dynamics remain challenging to interpret. In this study, we observed an increase in calcium activity in DG 24 hours after stimulation, despite the return of c-Fos expression to baseline. This discrepancy highlights the complexity of interpreting activity-dependent markers and suggests that c-Fos and calcium imaging may capture distinct aspects of neural activity. While c-Fos reflects overall neuronal activation involving both excitatory and inhibitory populations, calcium imaging primarily primarily measures fluctuations in intracellular calcium, which may be influenced by changes in neuronal excitability and synaptic activity rather than solely reflecting evoked responses. The observed increase in calcium activity, despite stable c-Fos expression, suggests that spontaneous firing rate homeostasis in the DG was maintained, indicating a possible dissociation between local synaptic potentiation and network-wide activity regulation. This preservation of homeostasis is critical for maintaining network stability and may represent an adaptive mechanism that prevents overstimulation while permitting targeted plastic changes.

In contrast to DG, the hilus exhibited an increase in calcium activity at both 3 and 24 hours post-stimulation. This region, which contains mossy cells and inhibitory interneurons, may play a key role in modulating local network dynamics through reciprocal interactions with granule cells. The elevated activity observed in the hilus could reflect enhanced excitatory input from granule cells, alongside compensatory inhibitory feedback to stabilize the overall network function. These findings highlight region-specific differences in iTBS-induced plasticity and underscore the importance of inhibitory circuits in shaping the network’s response to stimulation [42, 43].

The biphasic nature of the functional changes and spine remodeling observed in this study may reflect distinct phases of network adaptation to iTBS-induced synaptic potentiation. The initial phase, characterized by rapid spine enlargement and increased mEPSC amplitudes and frequencies, suggests immediate activity-dependent potentiation of specific synapses. This is likely accompanied by homeostatic mechanisms that stabilize spontaneous firing rates and prevent network hyperexcitability. The second phase of spine enlargement at 24 hours may represent secondary remodeling processes driven by network-level integration of the initial potentiation. These delayed structural changes could involve complex interactions between the excitatory and inhibitory circuits, resulting in optimized connectivity and long-term stabilization of the induced plasticity. Together, these findings emphasize the dynamic interaction between functional and structural plasticity, highlighting the role of temporal and spatial coordination in adaptation to stimulation-induced perturbations.

Notably, the introduction of multiple iTBS-rTMS sessions, as implemented in FDA-approved protocols, may interact with the temporal dynamics of the second phase of structural and functional remodeling observed in this study. By inducing additional waves of Hebbian and homeostatic plasticity during this phase, multi-session rTMS protocols may disrupt or amplify the secondary remodeling processes. This could result in a cascade of alternating Hebbian and homeostatic plasticity events, ultimately leading to profound structural and functional reorganization at both the microcircuit and the network levels. These overlapping waves of plasticity may account for the clinical efficacy of multi-session rTMS in promoting long-lasting therapeutic effects by leveraging the brain’s capacity for dynamic and adaptive remodeling.

## Supportive Information

### Funding

AV: National Institutes of Health, USA (NIH; 1R01NS109498) and by the Federal Ministry of Education and Research, Germany (BMBF, 01GQ1804A).

## Acknowledgments

The authors thank Dr. Christos Galanis and Dr. Dimitrios Kleidonas for the introduction and instructions on preparing OTCs and performing magnetic stimulation on OCTs. Useful discussions from Sigrun Nestel are greatly appreciated. The authors also thank the excellent lab support from technical assistants Birgit Egle and Monika Paetzold. In interpreting the experimental results, we acknowledge the useful inputs from Dr. Christian Tetzlaff and Dr. Jannik Luboeinski. The authors express sincere thanks to Pia Kruse, Dr. Matthias Kirsch, and Hanna Hemeling for exploratory experiments and analysis that we decided not to include in the current study to broaden the scope.

## Author contributions

HL, AV conceived the project. HL, ML, AV designed the experiments. HL, LH, ML conducted the experiments. HL quantified and analyzed the data. SG helped with the computer vision algorithm. HL prepared the figures and wrote the manuscript. All authors contributed to reviewing and proofreading.

